# MDR1 promotes CD8 T cell persistence in tumors and protects against cytotoxic chemotherapy

**DOI:** 10.1101/2025.09.28.679049

**Authors:** Lincoln A. Brown, Megan M. Erwin, Natalie R. Favret, Claudia N. McDavid, Jessica J. Roetman, Zachary D. Ewell, Melissa M. Wolf, Kristen A. Murray, Jack E. Smithwick, Marti A. Goemann, Mary Philip

## Abstract

Multidrug transporters, including multidrug resistance-1 (MDR1), are recognized chiefly for effluxing chemotherapeutic drugs out of tumor cells. However, they are also expressed in many normal cells and tissues, including lymphocytes, but their physiological role is less well-understood. Here, we investigated the role of MDR1 in tumor-specific CD8 T cells (TST), which are critical in antitumor immunity and key targets of immunotherapies. Using a clinically-relevant genetic liver cancer mouse model, we investigated the efflux dynamics of TST as they underwent activation, proliferation, and differentiation to dysfunctional states in tumor-bearing hosts. Surprisingly, we found that late-stage/terminally dysfunctional TST had the highest efflux capacity in both murine and human liver tumors. TST upregulated transcription of *Abcb1a*, encoding MDR1. We used CRISPR/Cas9 to generate MDR1-deficient TST, which persisted poorly in tumor-bearing mice as compared to MDR1-sufficient TST. MDR1 expression improved TST viability and reduced reactive oxygen species accumulation. Loss of MDR1 made T cells more susceptible to cytotoxic chemotherapy-induced cell death. Our findings demonstrate a role for MDR1 in regulating TST persistence and oxidative stress, with implications for antitumor T cell therapies in patients and immune regulation following cytotoxic chemotherapy.

## INTRODUCTION

ATP-binding cassette (ABC) transporters are membrane-associated transporters that import or export diverse substrates from cells^1^. A subset of ABC transporters, termed multidrug transporters, are known to efflux chemotherapeutic drugs out of cancer cells. Multidrug transporters include multidrug resistance-1 (MDR1, also known as P-glycoprotein; encoded by *ABCB1* in humans and *Abcb1a* and *Abcb1b* in mice), breast cancer resistance protein (BCRP; encoded by *ABCG2/Abcg2*), and multidrug resistance protein 1 (MRP1, encoded by *ABCC1/Abcc1*). These transporters are also present in many normal cell types, including immune cells, although their physiological roles are less understood^2^.

The expression of multidrug transporters varies across the hematopoietic system. MDR1 is detectable in some bone marrow precursors and most mature immune cells^3^. Side population cells, a subset of hematopoietic cells enriched for stem cells, are defined by their high capacity to efflux Hoechst dyes, mediated primarily by BCRP^4^. This increased efflux capacity has been suggested to protect hematopoietic stem cells from xenobiotics and promote survival following cytotoxic chemotherapy exposure. MDR1 is upregulated in lymphocytes in the gut and has been shown to protect ileal CD4 T cells from bile acid-driven stress; loss of MDR1 leads to pathogenic Th1 and Th17 T cells^5^. MDR1 is constitutively expressed in NK and CD8 T cells and is upregulated in select effector and memory subsets^3,6^. These findings establish a role for multidrug transporters in the function and homeostasis of T cells, yet little is known about their contribution to tumor-specific CD8 T cells (TST).

T cells are important for antitumor responses in patients and are central to cancer immunotherapies, including immune checkpoint blockade (ICB) and chimeric antigen receptor (CAR) T cell therapies^7^. To understand the role of multidrug transporters in TST, we utilized our genetic mouse model of liver cancer, in which adoptively transferred TST initially differentiate to an ‘early dysfunctional’ state (PD1^hi^ CD39^lo^ TCF1^hi^ TOX^hi^) and then to a ‘late dysfunctional’ state (PD1^hi^ CD39^hi^ TCF1^lo^ TOX^hi^), driven by persistent tumor antigen stimulation and epigenetically encoded^8,9^. We found that late dysfunctional TST display strikingly high efflux capacity, mediated by MDR1. Loss of MDR1 resulted in impaired TST persistence and an increased accumulation of mitochondrial reactive oxygen species (ROS). We demonstrated that MDR1 protects T cells exposed to cytotoxic chemotherapy. Taken together, our findings provide new insights into the role of MDR1 in T cell fitness in tumors, which have implications for understanding T cell responses to chemo- and immunotherapy and offer a potential therapeutic target to modulate antitumor T cell responses in patients.

## MATERIALS AND METHODS

### Mice

SV40 large T antigen (TAG)-specific CD8 T cell (TCR_TAG_) transgenic mice (Strain C57BL/6J Thy1.1 mice (Strain 000406) were purchased from The Jackson Laboratory. TCR_TAG_ mice were crossed to Thy1.1 mice to generate TCR_TAG_;Thy1.1^+^ mice. Cas9 mice were crossed to TCR_TAG_;Thy1.1 mice to generate Cas9;TCR_TAG_;Thy1.1^+^ mice. AST ((Albumin-floxStop-SV40 large T antigen(TAG))^10^ mice were crossed to Cre-ER^T2^ mice to generate AST;Cre-ER^T2^. Both female and male mice were used for studies. Mice were age- and sex-matched and between 6-11 weeks old when used for experiments. T cell donor mice were between 6-11 weeks of age and sex-matched to recipient male and female C57BL/6 and AST;Cre-ER^T2^ recipients. All mice were bred and housed in the animal facility at Vanderbilt University Medical Center (VUMC). All animal experiments were performed in compliance with VUMC Institutional Animal Care and Use Committee (IACUC) regulations and in accordance with approved VUMC IACUC protocol M1700166.

### Adoptive T cell transfer in tumor and acute infection models

To initiate tumorigenesis, AST;Cre-ER^T2^ mice were treated intraperitoneally with 1 mg tamoxifen (TAM) 1-2 days prior to adoptive T cell transfer. For the transfer of naive TCR_TAG_ T cells into AST;Cre-ER^T2^ mice, 1-2x10^6^ CD8 splenocytes from TCR_TAG_;Thy1.1^+^ transgenic mice were adoptively transferred into AST;Cre-ER^T2^ (Thy1.2^+^) mice. For the transfer of MDR1-deficient TCR_TAG_ (MDR1KO) or control TCR_TAG_ (ntc) into AST;Cre-ER^T2^ mice, 1x10^6^ TCR_TAG_ were adoptively transferred into AST;Cre-ER^T2^ mice within 6 hours of transfection (described below). For the generation of effector and memory TCR_TAG_, 2x10^5^ CD8 splenocytes from TCR_TAG_;Thy1.1^+^ transgenic mice were adoptively transferred into C57BL/6 (Thy1.2^+^) mice; within 2 hours mice were inoculated intravenously with 5-9x10^6^ CFU of *Listeria monocytogenes* (Lm) Δ*actA* Δ*inlB* strain^11^ expressing the TAG-I epitope (SAINNYAQKL, SV40 large T antigen 206–215).

### Cell isolation for subsequent analyses

Spleens from B6 or AST;Cre-ER^T2^ experimental mice were mechanically disrupted with the back of 3 mL syringe and filtered through a 70 mm strainer into ACK buffer. Cells were washed once and resuspended in cold complete RPMI (cRPMI) 1640 supplemented with 10% heat-inactivated FBS, 1 mM sodium pyruvate (Corning), 10 mM Hepes (Corning), 1X nonessential amino acids (Corning), 1X penicillin-streptomycin-glutamine (Thermo Fisher), and 55 µM 2-mercaptoethanol (Thermo Fisher) (cRPMI). Liver tissue was mechanically disrupted using a 150 mm metal mesh and glass pestle in ice-cold 2% FBS/PBS and passed through a 70 mm strainer. Liver homogenate was centrifuged at 400g for 5 min at 4°C and supernatant discarded. The liver pellet was resuspended in 15 mL of 2% FBS/PBS buffer containing 500 U heparin, mixed with 10 mL of Percoll (GE) by inversion, and centrifuged at 500g for 10 min at 4°C. Supernatant was discarded and the pellet was RBC lysed in ACK buffer and resuspended in cRPMI for downstream applications.

### AST1825 cell line

A tumor-bearing liver was isolated from a TAM-treated AST;Cre-ER^T2^ mouse, washed with PBS, and sterilely-minced into small pieces in 5 mL 2% FBS/PBS. An additional 5 mL of Liver Digest Medium (collagenase-dispase; Gibco) was added and the tissue digest incubated for 30 min at 37℃. The digested liver suspension was filtered through sterile medical gauze and centrifuged at 1000 rpm for 5 min at 4°C. The pellet was resuspended in 10 mL 2% FBS/PBS and filtered sequentially through 70 µm filter and 45 µm filters, centrifuged, and the supernatant discarded. The cell pellet was RBC lysed in ACK buffer for 5-10 min at room temperature and centrifuged at 1000 rpm for 5 min at 4°C. The resulting single-cell hepatocyte/hepatocellular carcinoma suspension were cultured in DMEM supplemented with 10% heat-inactivated FBS and 1X penicillin-streptomycin-glutamine (Thermo Fisher) (cDMEM) at 37℃ for 2 weeks and split at 1:5 ratio to establish the line. The established cell line was cryopreserved in FBS supplemented with 10% DMSO and stored in liquid N_2_. For use in experiments, cells were thawed and cultured in cDMEM. When cells reached 80% confluency, adherent cells were digested in trypsin-EDTA (0.25%; Thermo Fisher), washed with DMEM, and split at a 1:10 ratio.

### T cell culture

All T cell cultures were performed in a 37℃ 5% CO2 humidified incubator. To activate T cells in vitro, TCR_TAG_ were enriched from splenocytes using EasySep^TM^ Mouse CD8 T Cell Isolation Kit (STEMCELL Technologies) and resuspended in cRPMI at 0.5-1x10^6^ cells/mL with anti-CD3 (1 µg/mL; Tonbo), anti-CD28 (0.5 µg/mL; Tonbo), and 50 U/mL recombinant human IL-2 (rhIL-2; BRB Preclinical Biologics Repository). After 48 hours, media was replaced with cRPMI supplemented with 50 U/mL rhIL-2. Doxorubicin (100 nM; Sagent) was added to culture media where indicated. For T cell-AST1825 cell line coculture experiments, TCR_TAG_ were resuspended at 0.5x10^6^ cells/mL in cRPMI with 50 U/mL rhIL-2 and 10 ng/mL recombinant human IL-15 (rhIL-15; BRB Preclinical Biologics Repository) and plated in flat bottom plates on top of adherent AST1825 cells that had been seeded 1-2 days prior. Every 24 hours, cells were transferred to new wells with adherent AST1825 cells that had been seeded 1-2 days prior. Every 48 hours, media was replaced with fresh media. After 5-6 days in culture, media was supplemented with 10 ng/mL recombinant human IL-7 (rhIL-7; BRB Preclinical Biologics Repository) and 10 ng/mL rhIL-15.

### Human samples

Deidentified liver tumor samples from patients at VUMC were supplied by the Cooperative Human Tissue Network (CHTN) according to approved protocols (VUMC IRB #231173). On the day of collection, human liver tumor samples were mechanically disrupted and lymphocytes enriched as described above for mouse liver tumors. If analysis was not performed on day of collection, cells were cryopreserved in Bambanker serum-free cryopreservation medium (Nippon Genetics) and stored in liquid nitrogen. For analysis, cryopreserved cells were thawed by swirling in 37℃ water bath. Thawed cell suspension was added to prewarmed complete medium, centrifuged at 400g for 5 min, and resuspended in medium for downstream applications.

### Flow cytometry

Surface staining was performed at 4℃ for 20-30 minutes. Intracellular transcription factor staining was performed with the Foxp3/Transcription Factor Staining Buffer Kit (Tonbo) per manufacturer’s instructions. Dead cell staining was performed using Ghost Dye Red 780 (Cytek) according to manufacturer instructions. For MitoTracker Green (MTG) efflux, T cells in cRPMI were incubated for 30 minutes in a 37℃ 5% CO2 humidified incubator with 200 nM MitoTracker Green FM (Thermo Fisher M7514) or MitoSpy Green FM (Biolegend 424805) in the presence or absence of an efflux inhibitor. Verapamil (50 µM; Sigma-Aldrich), elacridar (100 nM; Sigma-Aldrich), or KO143 (1 µM; Selleck) were used as inhibitors where indicated. Efflux index was quantified as log_2_ fold change of MTG MFI in medium relative to inhibitor-treated sample. For DCV efflux, T cells in cRPMI were incubated for 60 minutes in a 37℃ 5% CO2 humidified incubator with DyeCycle Violet (10 μM; Thermo Fisher V35003) in the presence or absence of verapamil (50 µM; Sigma-Aldrich). For mitochondrial superoxide detection, T cells in cRPMI were incubated with MitoSOX Green (5 µM; Thermo Fisher M36006) in a 37℃ 5% CO2 humidified incubator for 30 minutes. After staining, cells were washed twice with cold 2% FBS/PBS before data acquisition on the cytometer. Anti-mouse antibodies used included: APC-CD279, PE-Cy7-CD39, BV510-CD90.1, APC-CD90.1, BV421-CD90.2, FITC-Ki-67, BV785-CD62L, FITC-TCR Vβ7 (from BioLegend); PerCP-Cy5.5-CD8a, APC-CD8a, PE-CD8a (from Tonbo); and PerCP-eFluor710-CD279 (from eBioscience). Anti-human antibodies used included: BV421-CD8, BV711-CD4, BV785-CD45RO, PE-Cy7-CD279, APC-CD39 (from BioLegend); and rF710-CD45RA (from Tonbo). All flow analysis was performed on the Attune NXT Acoustic Focusing Cytometer (ThermoFisher Scientific). Data was analyzed using FlowJo v.10.10.0 (Tree Star Inc.). The gating strategy used to identify TCR_TAG_ is shown in **Supplemental Fig. 1A** and was used in all adoptive transfer and *in vitro* experiments, unless otherwise specified.

### Electroporation/Transfection

Splenocytes from naive Cas9;TCR_TAG_;Thy1.1^+^ transgenic mice were isolated as described above. Cas9;TCR_TAG_ were enriched from splenocytes using EasySep^TM^ Mouse CD8 T Cell Isolation Kit (STEMCELL Technologies), counted using a hemacytometer, spun down, and resuspended in Opti-Mem (Thermo Fisher). To a 100 μL volume suspension of T cells (5-7.5x10^6^ cells) was added sgRNA targeting *Abcb1a* (GAAGACAGAUACACAAGAUC) and *Abcb1b* (CCAAACACCAGCAUCAAGAG) or a non-targeting control sgRNA (GACAUUUCUUUCCCCACUGG) (0.4 nmol) (Synthego) in a 2mm sterile cuvette (Bulldog Bio). Cells were electroporated using NEPA21 Electro-Kinetic Transfection System (Bulldog Bio) with parameters: Poring Pulse: 250 V, 2 ms length, 50 ms interval, 2 pulses, 10% Decay Rate, + polarity; Transfer Pulse: 20 V, 50 ms length, 50 ms interval, 5 pulses, 40% Decay Rate, +/- Polarity. Immediately after electroporation, pre-warmed cRPMI containing 100ng/ml of rhIL-7 was added to the cuvette, and the cuvette was incubated at 37℃ for 15 minutes. Cells were then gently mixed by pipetting and added to a flat-bottom 24-well plate. Additional warmed cRPMI was used to wash cuvette and added to wells. After 1-2 hours resting, cells were used for *in vitro* or *in vivo* experiments.

### Statistical analyses

No data were excluded from the analyses. The experiments were not randomized. The investigators were not blinded to allocation during experiments and outcome assessment. Statistical analyses on flow cytometric data were performed as described in the figure legends using Prism 10.5 software (GraphPad Software).

## RESULTS

### Efflux capacity increases with TST differentiation through dysfunctional states

The ability of cells to efflux fluorescent multidrug transporter substrates can be used to infer multidrug transporter expression patterns^12^. We assessed the ability of naive CD8 T cells to efflux the mitochondrial labeling dye MitoTracker Green (MTG), a known substrate of multidrug transporters^13^. Naive transgenic CD8 T cells specific for the SV40 large T antigen (TAG) epitope-I (TCR_TAG_)^14^ were incubated with MTG in the presence of the non-specific ABC transporter inhibitor, verapamil, or in medium as control, and then MTG fluorescence was assessed by flow cytometry. Comparing MTG fluorescence in medium versus verapamil-treated cells allowed for the assessment of efflux activity, with naive CD8 T cells showed modest MTG efflux (**Fig. 1A**).

**Figure 1.**
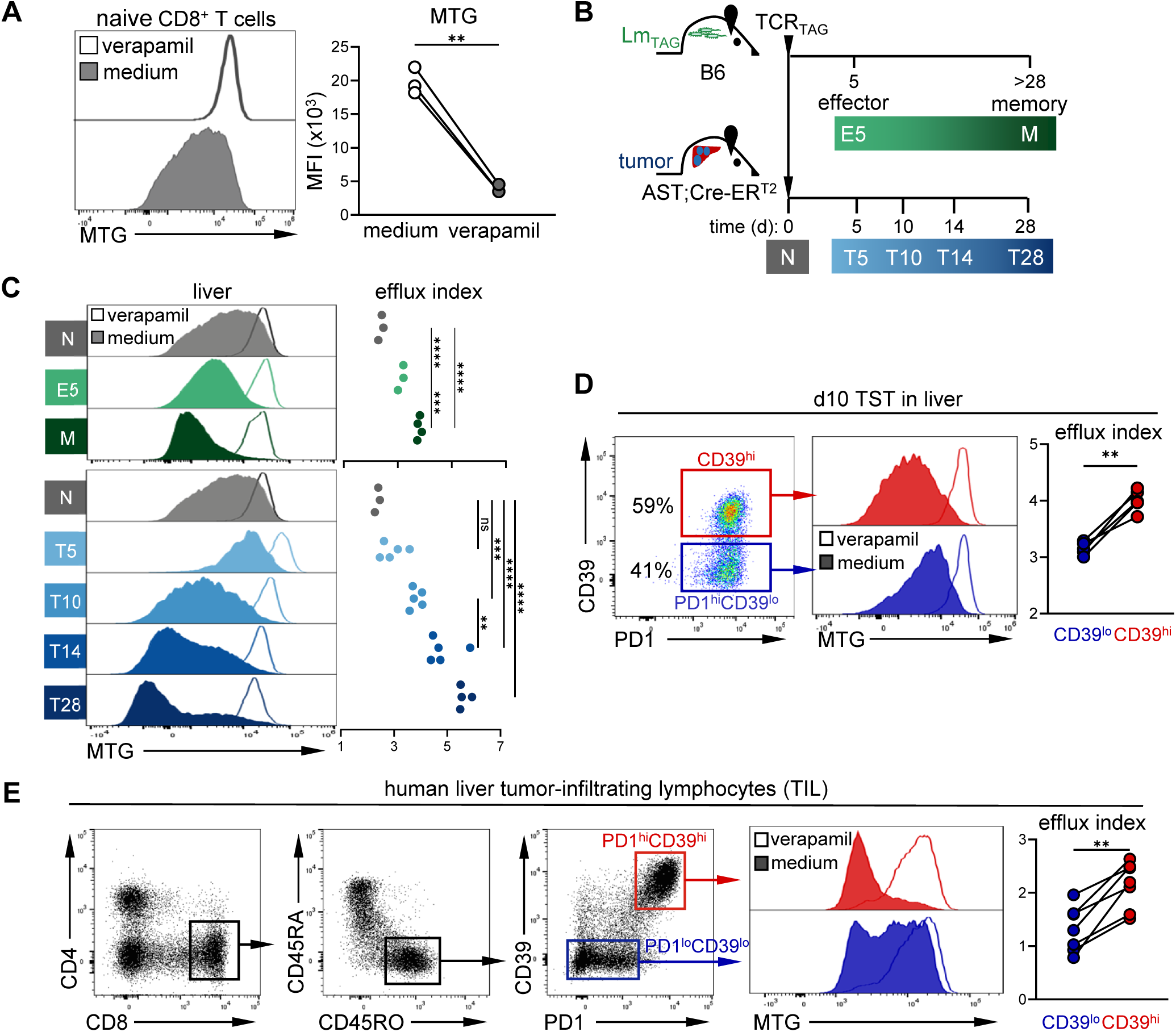
Efflux capacity increases with TST differentiation through dysfunctional states. **(A)** Mitotracker green (MTG) efflux in naive TCR_TAG_ CD8 T cells. Representative histograms show MTG fluorescence in TCR_TAG_ splenocytes incubated at 37°C with MTG in the medium alone (unfilled) or medium with verapamil (filled). Summary plot shows MTG MFI as paired samples for each mouse (+/- verapamil). \*\**P* ≤ 0.01; paired two-tailed Student’s t-test, n = 3, representative of three independent experiments. **(B)** Experimental scheme: naive TCR_TAG_ (Thy1.1^+^) were adoptively transferred into TAM-treated AST;Cre-ER^T2^ (Thy1.2^+^) or Lm_TAG_-infected C57BL/6 (Thy1.2^+^) mice. The same congenic transfer strategy is used for subsequent experiments and figures. Lymphocytes were re-isolated and analyzed by flow cytometry at 5, 10, 14, and 28 days post-transfer in tumor-bearing mice (T5-T28) or 5 and 30+ days post-transfer for Lm_TAG_ infected mice (E5 and M). **(C)** MTG fluorescence of TCR_TAG_ isolated from livers of infected or tumor-bearing mice as in (B). Summary plot shows the efflux index, with each dot showing a biological replicate, calculated as the log_2_ fold change of MTG MFI in medium relative to verapamil-treated sample. \*\**P*≤ 0.01, \*\*\**P*≤ 0.001, \*\*\*\**P* ≤ 0.0001; one-way ANOVA followed by Tukey’s multiple comparisons test; n = 3-5 per timepoint, representative of three independent experiments. **(D)** Efflux capacity of PD1^hi^CD39^lo^ and PD1^hi^CD39^hi^ TCR_TAG_ isolated from liver tumors 10d post-transfer. Left, gating strategy for PD1^hi^CD39^lo^ and PD1^hi^CD39^hi^ cells. Representative histogram showing MTG fluorescence. Right, summary plot of efflux index. \*\**P*≤ 0.01; paired two-tailed Student’s t-test, n = 5, representative of three independent experiments. **(E)** Efflux capacity of human tumor-infiltrating lymphocytes (TIL) isolated from fresh human hepatocellular carcinoma samples. Left, gating strategy for CD8CD45RO^+^PD1^hi^CD39^hi^ (tumor-reactive; red) and CD8CD45RO^+^PD1^lo^CD39^lo^ (bystander; blue) TIL. Right, TIL MTG fluorescence and summary plot of efflux index. \*\**P* ≤ 0.01; paired two-tailed Student’s t-test, n = 7 individual patient samples.

We next used our previously developed tamoxifen (TAM)-inducible genetic mouse model of liver cancer (AST;Cre-ER^T2^), in which liver tumors express the SV40 large T antigen (TAG), to measure efflux dynamics in TST over time. Naive transgenic TAG-specific CD8 T cells (TCR_TAG_) were adoptively transferred into TAM-treated AST;Cre-ER^T2^ mice or TAG epitope I-expressing *Listeria monocytogenes* (LM_TAG_)-infected C57BL/6 (B6) mice for comparison in the setting of acute infection. We then harvested and analyzed TCR_TAG_ from spleens and livers of the mice at several timepoints post-transfer (**Fig. 1B**). The flow cytometry gating strategy for TCR_TAG_ is shown in **Supplemental Fig. 1A**. As previously observed^3^, efflux capacity increased with activation during acute infection, with memory T cells having higher efflux capacity than effector and naive CD8 T cells (**Fig. 1C**). CD8 T cell efflux capacity also increased with activation in tumor-bearing hosts both in liver tumors (day 5; d5) (**Fig. 1C**) and in the spleen (**Supplemental Fig. 1B**). Interestingly, with increasing duration of tumor/antigen exposure, TST efflux capacity increased further in both livers and spleens, with d28 TST having the highest efflux (**Fig. 1C**). To ensure that these findings were not specific to MTG, we assessed efflux capacity in TST at d5 and 28 using DyeCycle Violet (DCV), a DNA dye and ABC transporter substrate that has been used to identify highly-effluxing cell populations^15^. TST at d28 showed greater ability to efflux DCV than d5 (**Supplemental Fig. 1C**).

We previously showed that TST differentiate to phenotypically and epigenetically distinct ‘early’ and ‘late’ dysfunctional states, driven by persistent tumor antigen exposure^8^. As we observed that TST efflux capacity increased over time, we asked if efflux capacity correlated with markers of T cell dysfunctional states. TST were largely PD1^hi^ CD39^lo^ at d5 (early dysfunctional) and PD1^hi^ CD39^hi^ (late dysfunctional) by d14+ (**Supplemental Fig. 1D**). At d10, an intermediate timepoint, there was heterogeneity, with a sizable population of both PD1^hi^ CD39^lo^ and PD1^hi^ CD39^hi^ TST within tumor (**Supplemental Fig. 1D**). We analyzed TST at the d10 intermediate time point and found that late/PD1^hi^ CD39^hi^ TST had a much higher efflux capacity than early/PD1^hi^ CD39^lo^ TST from the same tumor (**Fig. 1D**).

To determine whether increased efflux capacity was also a feature in human cancers, we examined tumor-infiltrating lymphocytes (TIL) isolated from resected tumors from patients with hepatocellular carcinoma (**Supplemental Table 1**). Gating on non-naive CD45RO^+^ CD8 T cells in liver tumors, we found that the TIL population was heterogeneous, with both PD1^lo^ CD39^lo^ TIL and PD1^hi^ CD39^hi^ TIL (**Fig. 1E**). Increased human TIL expression of PD1 and CD39 is associated with terminal exhaustion^16,17^ and expression of the dysfunction/exhaustion-associated transcription factor TOX^9^. Accordingly, more dysfunctional-appearing CD45RO^+^ PD1^hi^ CD39^hi^ TIL showed higher efflux capacity than CD45RO^+^ PD1^lo^ CD39^lo^ TIL (**Fig. 1E**), consistent with findings from our mouse model.

### MDR1 is upregulated in late TST and mediates TST efflux

We next sought to determine which multidrug efflux transporters were responsible for increased efflux capacity in TST. Analyzing our published transcriptional data^8^, we observed that *Abcb1a*, encoding MDR1A, was upregulated in late dysfunctional TST (d14+) as compared to other efflux protein-encoding genes (**Fig. 2A**). To test if MDR1 is the key player mediating TST efflux, we incubated early (d5) and late (d21) TST (**Fig. 2B**) with MTG in the presence or absence of the efflux inhibitors elacridar (MDR1-specific), KO143 (BCRP-specific), or verapamil (non-specific). We observed maximal efflux inhibition with elacridar, while KO143 had little effect, suggesting that MDR1 is chiefly responsible for TST efflux capacity (**Fig. 2C**). Elacridar also fully inhibited MTG efflux in effector and memory TCR_TAG_, indicating that MDR1 is the primary transporter responsible for MTG efflux in these T cell subsets as well (**Supplemental Fig. 2A, B**).

**Figure 2.**
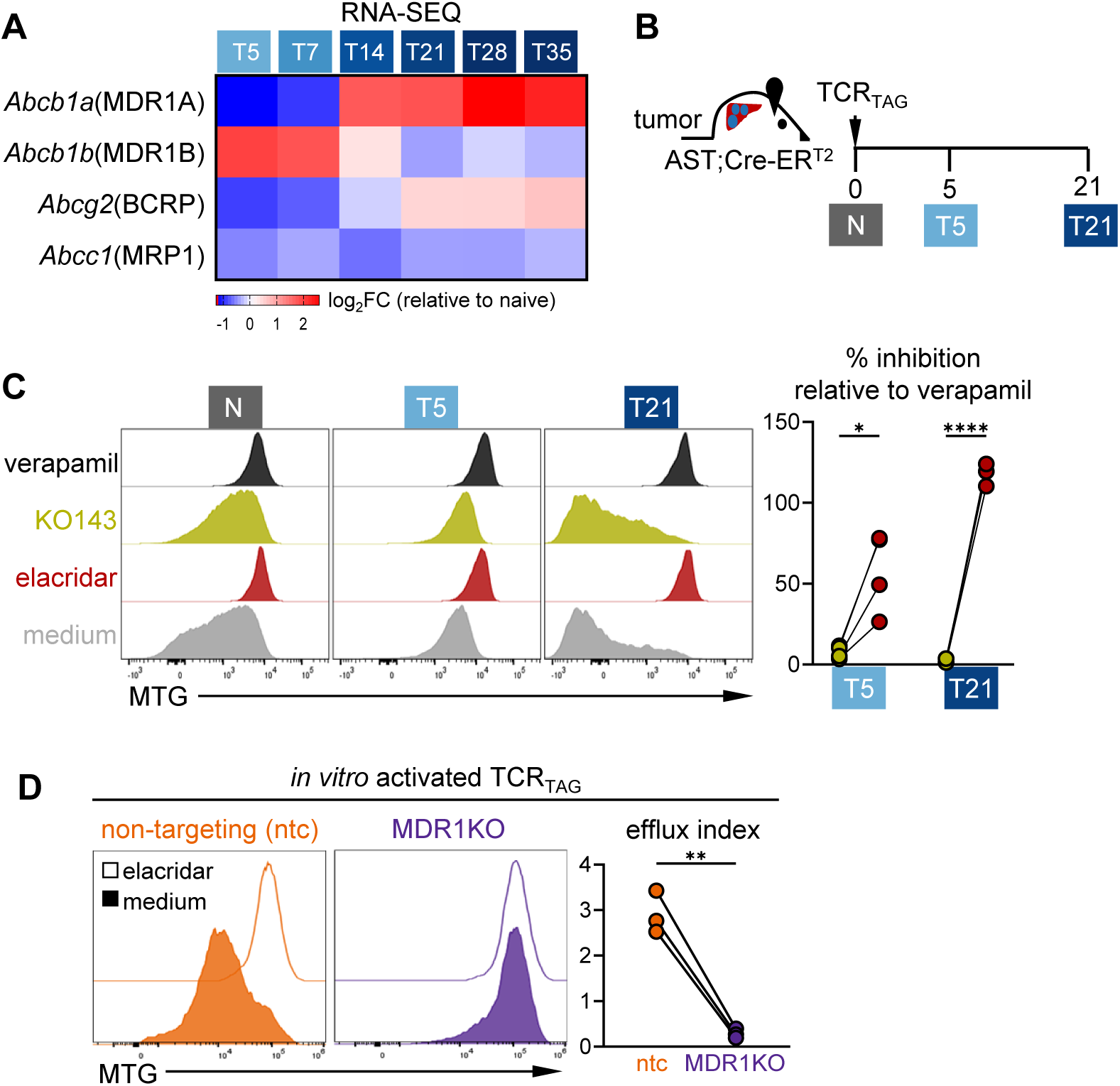
MDR1 is upregulated in late TST and mediates TST efflux. **(A)** Heatmap showing TST multidrug transporter gene expression, determined by RNA-SEQ (GSE89307) (log_2_ fold change compared to naive) for TCR_TAG_ at 5, 14, 21, 28, and 35d post-transfer as in Fig. 1B. Each gene is shown together with the transporter protein it encodes. n = 3 per timepoint. **(B)** Experimental scheme: naive TCR_TAG_ were adoptively transferred into TAM-treated AST;Cre-ER^T2^mice and lymphocytes re-isolated from livers 5 and 21d post-transfer. **(C)** MTG efflux of TCR_TAG_ incubated with indicated inhibitors. Summary plot shows % inhibition calculated from MTG MFI compared to verapamil-treated control. \**P* ≤ 0.05, \*\*\*\**P* ≤ 0.0001; paired Student’s t-test with Holm-Šídák correction for multiple comparisons, n = 4 per group. **(D)** Naive TCR_TAG_ were transfected with sgRNAs targeting *Abcb1a* and *Abcb1b* (MDR1KO; purple) or a non-targeting control sgRNA (ntc; orange), expanded *in vitro* with anti-CD3, anti-CD28, and IL-2, and MTG efflux assessed after 5 days. Representative histograms show MTG fluorescence and summary plot shows efflux index. \*\**P* ≤ 0.01; paired two-tailed Student’s t-test, n = 3, pooled from three independent experiments.

To target MDR1 expression more specifically, we used CRISPR/Cas9 to knock out the genes encoding MDR1 in TCR_TAG_. We transfected TCR_TAG_ with sgRNAs targeting *Abcb1a* and *Abcb1b* (MDR1KO) and assayed efflux capacity. MDR1KO TCR_TAG_ had minimal efflux capacity as compared to cells receiving a <u>n</u>on-targeting <u>c</u>ontrol sgRNA (ntc) and caused a similar degree of efflux impairment as the MDR1 inhibitor elacridar (**Fig. 2D**).

### MDR1-deficient TST persist poorly in tumors and accumulate mitochondrial ROS

To understand the role of MDR1 in TST, we adoptively transferred MDR1KO or ntc TCR_TAG_ into TAM-treated AST;Cre-ER^T2^ mice. After 20 days, TCR_TAG_ were harvested from livers and spleens (**Fig. 3A**). MDR1KO TST efflux capacity remained impaired 20 days after adoptive transfer (**Fig. 3B**). We observed markedly reduced numbers of MDR1KO TST in both the livers and spleens of AST;Cre-ER^T2^ mice, indicating that MDR1 supports TST persistence (**Fig. 3C**). MDR1 deficiency did not affect the immunophenotype/differentiation to dysfunctional states as assessed by PD1/CD39 co-expression (**Fig. 3D**), nor did MDR1 loss impact proliferation as determined by Ki67 expression (**Fig. 3E**). Prior studies have shown that MDR1 reduces reactive oxygen species (ROS) accumulation and supports mitochondrial function in a number of cells and tissues, including colonic epithelium, oocytes, and effector CD8 T cells^3,18,19^. To assess mitochondrial ROS in TCR_TAG_, we stained cells with MitoSOX, a fluorogenic dye that selectively reacts with mitochondrial superoxide^20^. In line with previousstudies, we observed increased MitoSOX staining in MDR1KO TST, indicating increased accumulation of mitochondrial ROS (**Fig. 3F**).

**Figure 3.**
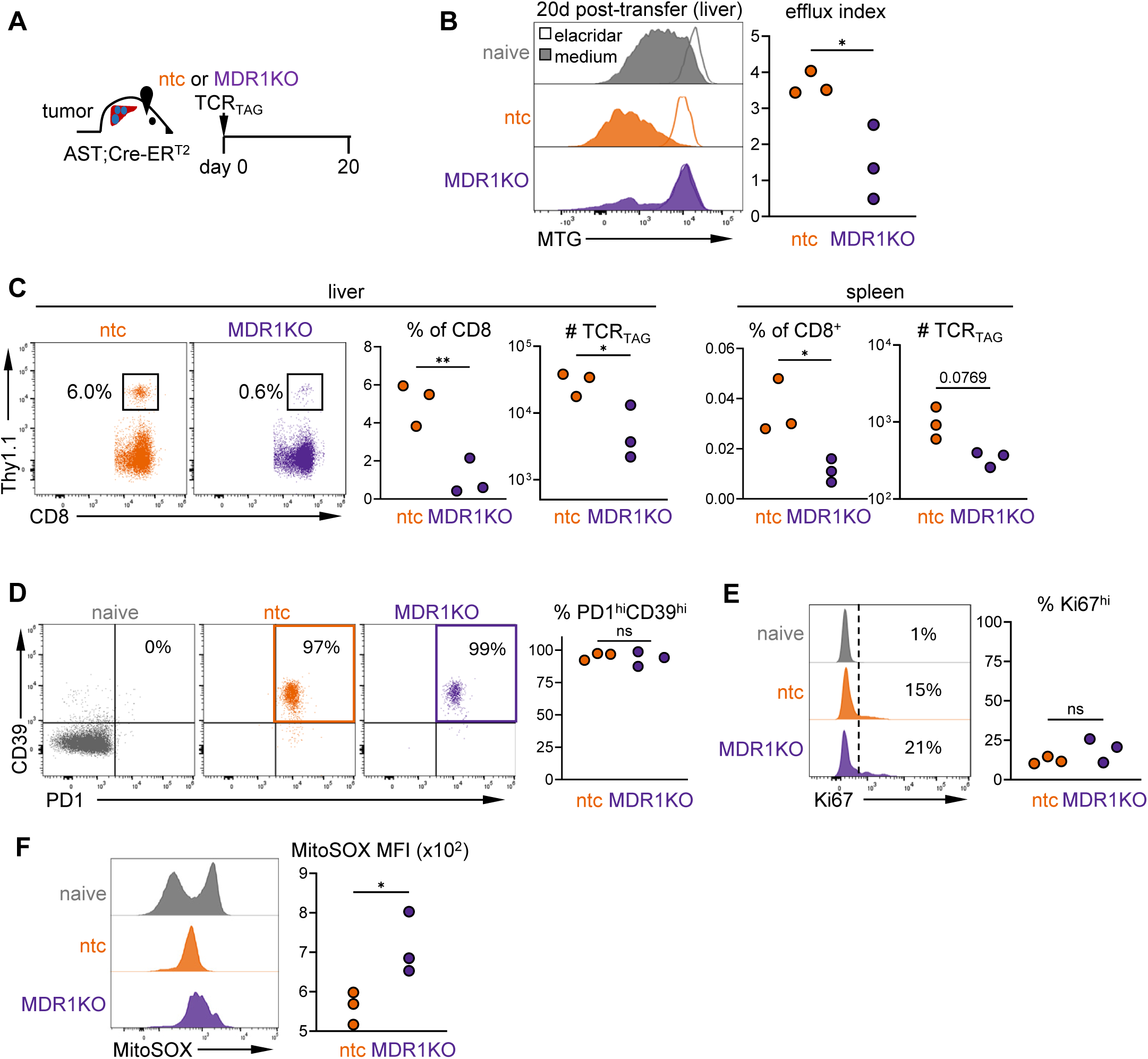
MDR1-deficient TST persist poorly and accumulate mitochondrial ROS. **(A)** Experimental scheme: naive MDR1-sufficient (ntc) or MDR1-deficient (MDR1KO) TCR_TAG_ were adoptively transferred into TAM-treated AST;Cre-ER^T2^ mice. Lymphocytes were re-isolated from livers and spleens and analyzed by flow cytometry at day 20 post-transfer. **(B)** Efflux capacity of ntc and MDR1KO. Summary plot shows efflux index. \**P* ≤ 0.05; two-tailed Student’s t-test, n = 3, representative of three independent experiments. **(C)** Flow plots showing ntc and MDR1KO TCR_TAG_ gated within the total CD8 T cells in the liver. Summary plots show TCR_TAG_ as % of live CD8 T cells and total TCR_TAG_ in the liver (left plots) and spleen (right). \**P* ≤ 0.05, \*\**P* ≤ 0.01; two-tailed Student’s t-test, n = 3 per group, representative of three independent experiments. **(D)** PD1 and CD39 expression of ntc and MDR1KO TCR_TAG_, gate set based on naive TCR_TAG_. ns = not significant; two-tailed Student’s t-test, n = 3, representative of three independent experiments. **(E)** Representative histogram of Ki67 expression, gate set based on naive TCR_TAG_. Summary plot shows % Ki67^hi^, ns = not significant; two-tailed Student’s t-test, n = 3, representative of three independent experiments. **(F)** MitoSOX staining of ntc and MDR1KO TCR_TAG_. Summary plot shows MitoSOX MFI, \**P* ≤ 0.05; two-tailed Student’s t-test, n = 3, representative of at least three independent experiments.

To further investigate the role of MDR1 in TST persistence, we used an *in vitro* model of chronic tumor antigen stimulation. TCR_TAG_ were cocultured and restimulated daily with a TAG-expressing liver cancer cell line (AST1825) generated from the liver of an AST;Cre-ER^T2^ mouse (**Fig. 4A**). Similar to what we observe in vivo in tumors, naive ntc and MDR1KO TCR_TAG_ cocultured with AST1825 cancer cells became activated, upregulated PD1 (**Fig. 4B**), and expanded over the first 7 days before then slowly declining in numbers (**Fig. 4C**). ntc TCR_TAG_ gained increased efflux capacity with time, similar to TST *in vivo*, while MDR1KO TCR_TAG_ continued to have impaired efflux (**Fig. 4D**). MDR1KO TCR_TAG_ showed decreased survival, particularly during the contraction phase (**Fig. 4C**). The loss of MDR1 did not appear to impact proliferation, as shown by a similar initial increase in cell numbers (**Fig. 4C**) and based on Ki67 expression (**Fig. 4E**). Instead, MDR1KO TCR_TAG_ underwent greater cell death (**Fig. 4F**) and accumulated more mitochondrial ROS (**Fig. 4G**), as we observed in tumors (**Fig. 3F**). Taken together, our findings support a role for MDR1 in regulating oxidative homeostasis and protecting TST from cell death, supporting long-term persistence.

**Figure 4.**
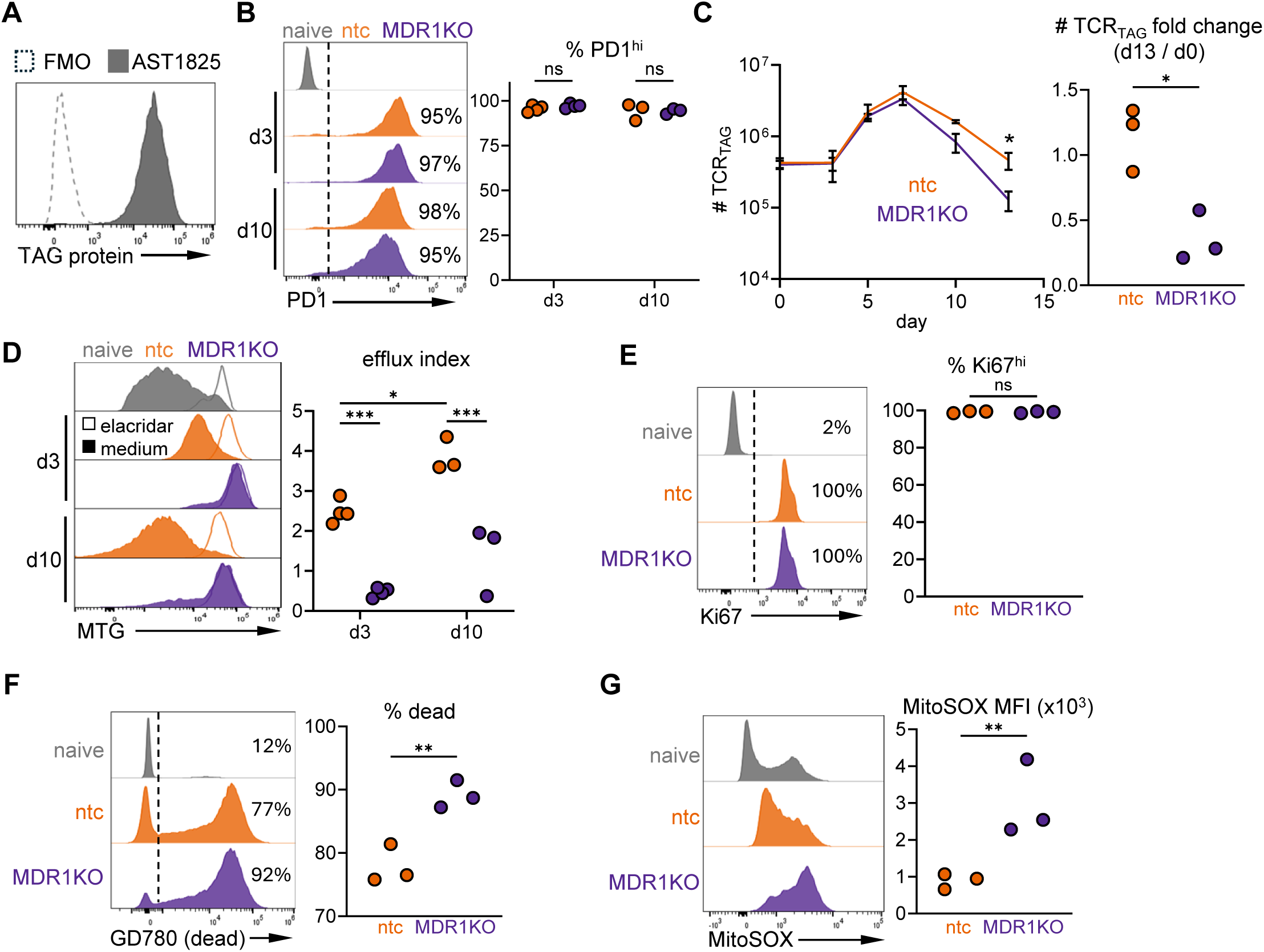
MDR1 promotes CD8 T cell viability during chronic tumor antigen stimulation. **(A)** Histogram showing SV40 large T Antigen (TAG) expression in AST1825 cell line. FMO (fluorescence minus one) shown as control. **(B)** Histogram showing ntc and MDR1KO TCR_TAG_ PD1 expression after 3 and 10 days (d) of coculture with AST1825, with gate set based on naive. Summary plot shows %PD1^hi^. ns = not significant; two-tailed Student’s t-test with Holm-Šídák correction for multiple comparisons, n = 3-4 biological replicates per timepoint, pooled from three independent experiments. **(C)** Left, number of ntc (orange) and MDR1KO (purple) TCR_TAG_ over time course of coculture with AST1825 cell line. Line plotted at mean with error bars showing ± SEM. \**P* ≤ 0.05; two-tailed lognormal t-test, n = 3-4 biological replicates, pooled from three independent experiments. Right: day 13 ntc and MDR1 KO TCR_TAG_ number relative to starting number, expressed as fold-change. n = 3 biological replicates, pooled from two independent experiments. \**P*≤ 0.05; two-tailed Student’s t-test. (**D**) Efflux capacity of ntc and MDR1KO TCR_TAG_ after 3 and 10d coculture with AST1825. Summary plot shows efflux index, \**P* ≤ 0.05, \*\*\**P* ≤ 0.001; one-way ANOVA followed by Tukey’s multiple comparisons test, n = 3-4 biological replicates per timepoint, pooled from three independent experiments. **(E)** Ki67 expression of ntc and MDR1KO TCR_TAG_ after 5 days culture with AST1825 with gate set based on naive TCR_TAG_. Summary plot shows %Ki-67^hi^. ns = not significant; two-tailed Student’s t-test, n = 3 biological replicates, pooled from two independent experiments. **(F)** Viability of ntc and MDR1KO TCR_TAG_ after 12d culture with AST1825. Representative histograms show Ghost Dye Red 780 (GD780) staining, with gate for dead cells set based on naive TCR_TAG_. Summary plot shows % dead. \*\**P* ≤ 0.01; two-tailed Student’s t-test, n = 3 biological replicates, pooled from two independent experiments. **(G)** MitoSOX staining of ntc and MDR1KO TCR_TAG_ after 13d culture with AST1825. Summary plot shows MitoSOX MFI. \*\**P* ≤ 0.01; two-tailed Student’s t-test, n = 3 biological replicates, pooled from two independent experiments.

### MDR1 protects T cells from cytotoxic chemotherapy

Cytotoxic chemotherapies used in cancer treatments can potentiate antitumor immunity and are increasingly being used in combination with immunotherapies^21^. Doxorubicin is an anthracycline-based chemotherapy used in many cancer types that has been observed to selectively enhance CD8 TIL responses^22^, and has been used in combination with ICB in breast cancer and sarcomas^23,24^. However, exactly how different cytotoxic chemotherapies interact with immune cells and immunotherapy has not been well-explored. Given that many chemotherapeutic agents, including doxorubicin, are substrates of MDR1 and known promoters of oxidative stress^25^, we asked how MDR1 expression impacts T cell susceptibility to cytotoxic chemotherapy.

We generated naive ntc or MDR1KO TCR_TAG,_ which we then activated *in vitro*. We exposed the activated T cells to doxorubicin at a therapeutic dose (100 nM)^26^ (**Fig. 5A**). Doxorubicin treatment led to an overall decrease in the number of TCR_TAG_, however the impact on MDR1KO was markedly increased as compared to ntc (**Fig. 5B**). MDR1KO TCR_TAG_ underwent cell death at a much higher rate (**Fig. 5C**), indicating that MDR1 protects T cells from chemotherapy-induced cell death. Our findings in tumor-specific CD8 T cells, together with those of other groups investigating efflux and MDR1 expression in other immune cell subsets, suggest that MDR1 efflux activity, through its impact on cell survival, could play an important role in shaping the immune landscape in patients receiving chemotherapy.

**Figure 5.**
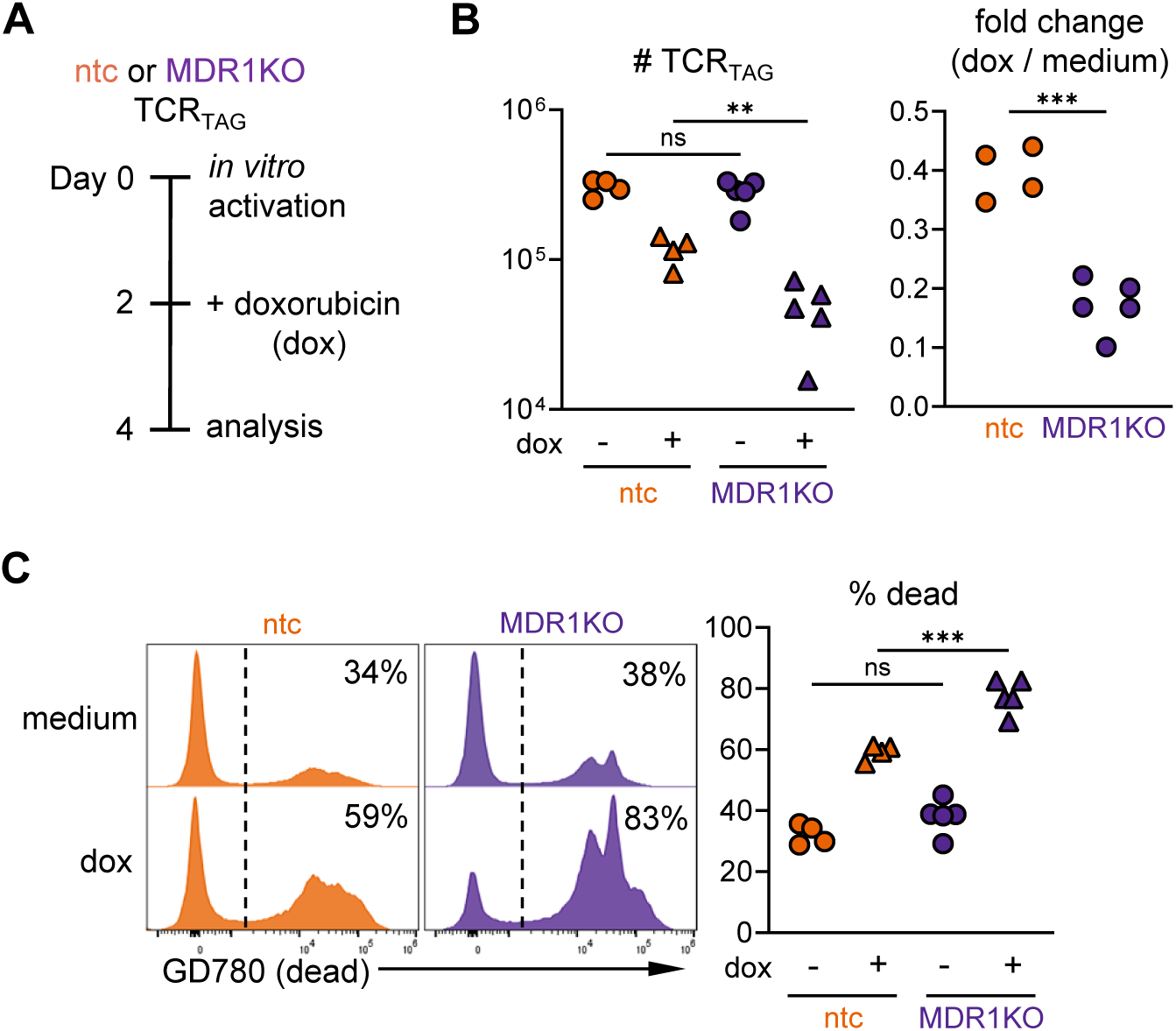
MDR1 protects CD8 T cells from cytotoxic chemotherapy. **(A)** Experimental scheme: Naive ntc and MDR1KO TCR_TAG_ were activated *in vitro* with anti-CD3, anti-CD28, and IL-2. On day 2, doxorubicin (dox) (100 nM) was added to the media. Cells were analyzed by flow cytometry on day 4. **(B)** Expansion of ntc (orange) and MDR1KO (purple) TCR_TAG_ in the presence or absence of dox. Left, number of TCR_TAG_. ns = not significant, \*\**P* ≤ 0.01; lognormal one-way ANOVA followed by Tukey’s multiple comparisons test; n = 4-5 biological replicates, pooled from three independent experiments. Right, fold change in number of dox-treated cells compared to untreated cells. \*\*\**P* ≤ 0.001; two-tailed Student’s t-test, n = 4-5 biological replicates, pooled from three independent experiments. **(C)** Viability of ntc and MDR1KO TCR_TAG_. Representative histograms show Ghost Dye Red 780 (GD780) staining with gate for dead cells set based on untreated TCR_TAG_. Summary plot shows % dead. ns = not significant, \*\*\**P* ≤ 0.001; one-way ANOVA followed by Tukey’s multiple comparisons test, n = 4-5 biological replicates.

## DISCUSSION

Here, we described MDR1-mediated efflux dynamics as CD8 T cells differentiated in the context of tumors (mice and patients with liver cancer) and during acute infection. Effector and memory T cells had higher efflux capacity than naive T cells, and TST in both murine and human tumors also had increased efflux capacity. Notably, TST exposed to tumor antigen over long periods and expressing markers of terminal dysfunction/exhaustion (PD1^hi^ CD39^hi^) expressed the highest level of MDR1 and exhibited the greatest efflux capacity. Using pharmacologic and genetic approaches, we showed that MDR1 was the key transporter mediating TST efflux. MDR1 was required for optimal TST persistence in an autochthonous liver cancer mouse model and an *in vitro* model of chronic tumor antigen stimulation. MDR1 deficiency did not impact T cell proliferation or alter T cell differentiation in tumors or during acute infection. Rather, MDR1 loss caused increased mitochondrial ROS in TST and decreased survival.

While MDR1 and other efflux transporters were initially studied for their ability to efflux drugs and xenobiotics, their physiologic role and substrates has been more difficult to parse. A conserved role for MDR1 is supported by the observation that MDR1 is widely expressed throughout the hematopoietic system^3^, and high efflux capacity is a feature of disparate immune subsets^2^. In CD8 T cells responding to acute infection, Chen *et al*. demonstrated that MDR1 promoted survival and mitochondrial fitness^3^. Our study presents a previously unappreciated role for MDR1 in long-term TST persistence and is supported by a recent study showing that deletion of the transporters SLCO3A1 and MDR1 in murine TST in the same liver cancer model led to reduced TST numbers, potentially due to increased oxidative stress caused by conjugated bile acids. Moreover, overexpression of SLCO3A1 improved TST persistence^27^. While this study suggested TST have low efflux capacity, the focus was primarily on early TST. Our work expands on these findings, showing that efflux capacity is dynamically regulated in TST over time. Notably, efflux capacity is highest in TST with longer (>14 days) exposure to tumor/tumor-bearing host or evidence of differentiation to the late dysfunctional state (PD1^hi^ CD39^hi^). We found increased TST efflux capacity in late dysfunctional TST in both the liver and spleen of tumor-bearing mice. Moreover, MDR1 deletion led to decreased survival of TST in both the liver and spleen, suggesting that MDR1 mitigates against factors besides bile acids or xenobiotics. A complete understanding of the endogenous substrates of MDR1 is still lacking.

Interestingly, increased MDR1 expression and efflux activity have been associated with the expression of the C-type lectin CD161 (KLRB1). Turtle *et al.* identified a self-renewing stem cell-like memory CD161^+^ CD8 T cell subset with high MDR1 expression that displayed the greatest ability to survive chemotherapy exposure^28^. CD161^+^ CD8 T cells are enriched in the gut and liver and have high efflux capacity, thought to support survival in tissues with high xenobiotic exposure^29^. High MDR1 expression is also recognized as a feature of CD161^+^ mucosal-associated invariant CD8 T cells (MAIT), which show marked resistance to xenobiotics and chemotherapy^30^. Among CD4 T cells, a self-renewing, viral-specific memory CD161^+^ subset that survives chemotherapy has been identified with high MDR1 expression^31^. Notably, we and others have shown that CD161 is upregulated in TIL in a variety of tumors and in dysfunctional chimeric antigen receptor T cells (CAR T)^8,32^. Indeed, a common phenotype and conserved transcriptional signature have been observed in CD161-expressing human T cells across various lineages, with key features including upregulation of *MDR1* and tissue-homing receptors^33,34^. These studies indicate that highly effluxing T cells share key features and suggest underlying shared regulation of MDR1 and CD161. One potential candidate is the Runt-related (RUNX) transcription factor RUNX3, a known regulator of T cell tissue residency^35^ which has been shown to maintain MDR1 expression in CD8 T cells^3^. While MDR1 upregulation may be mediated in part by RUNX3 and tissue residency programs, given that we observed increased TST efflux in secondary lymphoid organs/spleen, MDR1 is likely also regulated by other pathways.

One possibility is suggested by the fact that we observed a greater impact of MDR1 deletion on TST survival in response to long-term antigen stimulation, both *in vitro* and *in vivo*. TCR signaling is known to produce mitochondrial ROS, which are required for effective activation and expansion and play a role in activation-induced cell death^36,37^. However, mitochondrial ROS can cause cellular damage and drive T cell dysfunction, including through promoting telomeric damage^38–40^. MDR1 upregulation in late TST may therefore be an adaptation to chronic antigen stimulation. Chronic antigen stimulation and an immunosuppressive tumor microenvironment promote T cell dysfunction and poor persistence of endogenous and adoptively-transferred CD8 T cells^8,9,41^, limiting the efficacy of immunotherapies like ICB and CAR T. Understanding regulators of TST fitness is critical to improving and developing new immunotherapies. Augmenting MDR1 in adoptively-transferred CAR T or TCR-transduced T cells could reduce oxidative stress and improve persistence.

Alongside the potential role of MDR1 in protecting TST against mitochondrial stress induced by chronic TCR signaling, our findings have important clinical implications for MDR1’s role in effluxing chemotherapeutic agents. Chemotherapy is a cornerstone of cancer treatment and has direct cytotoxic effects on tumor cells as well as significant immunomodulatory properties. Interestingly, chemotherapy can promote antitumor immunity through several mechanisms, including enhancing the antigenicity of tumor cells and modulating the immune infiltrate, and it is increasingly being utilized in combination with ICB^21,42^. Chemotherapy selectively depletes immunosuppressive cells such as regulatory T cells and has been observed to increase the infiltration of CD8 T cells in tumors^43,44^. Among tumor-infiltrating CD8 T cell subsets, chemotherapy selectively expands TCF-1^+^ stem-like and precursor-exhausted TST^45^. We showed that MDR1 protects CD8 T cells from cytotoxic chemotherapy, potentially offering a mechanistic explanation for the previously observed enrichment of CD8 T cells following chemotherapy^22^. In line with this, a CXCR1^+^ CD8 T cell subset with high efflux capacity has been described in patients, which can withstand chemotherapy and is responsive to ICB^46^. Additionally, intratumoral MDR1-expressing CD4 T cell subsets were observed to be enriched after neoadjuvant chemotherapy in breast cancer, while T_reg_ cells, which lack MDR1, were depleted^47^. Taken together, these findings suggest that MDR1 regulates T cell responses and shapes the tumor immune infiltrate in patients receiving chemotherapy.

MDR1 inhibitors such as elacridar have been trialed as a strategy to promote chemo-sensitivity of tumor cells but have been largely unsuccessful clinically^48^. Our finding that MDR1 is required for optimal TST persistence may provide a possible explanation—increased cancer cell chemo sensitivity induced by MDR1 inhibition may be counteracted by impaired immune responses. However, this negative effect could be a potential benefit in the context of lymphodepletion given prior to adoptive T cell therapy, in which the goal is to clear endogenous lymphocytes, including dysfunctional TIL, and promote adoptive T cell engraftment^49^. Thus, in targeting MDR1, the impact on cancer cells as well as on the immune system must be carefully weighed depending on the clinical context.

In summary, our study examined the efflux dynamics of TST and identified late TST as a subset with high efflux capacity, mediated by MDR1. We show that MDR1 promotes TST persistence, reduces mitochondrial ROS accumulation, and protects T cells from cytotoxic chemotherapy, providing novel insight into the role of MDR1 in T cells. Our findings contribute to the understanding of the interplay between chemotherapy and the immune system and offer new potential avenues to modulate endogenous and adoptive T cell responses to improve anticancer therapies.

## ACKNOWLEDGEMENTS

We thank the Vanderbilt Division of Animal Care. We thank K. Wiles and the Cooperative Human Tissue Network (CHTN). We thank A. Schietinger and K.M. Hawley for technical assistance with the RNA electroporation protocol. This work was supported by the following funding sources: American Society of Hematology Medical Student Physician-Scientist Award (L.A.B.), NIH R37CA263614 (M.P.), NIH T32GM008554 (N.R.F.), and NIH T32AR059039 (M.M.E.).

## AUTHOR CONTRIBTIONS

(L.A.B.) and (M.P.) conceived and designed the study and analyzed and interpreted data. (L.A.B.) carried out experiments, assisted by (M.M.E.), (N.R.F.), (C.N.M.), (Z.D.E.), (M.M.W.), (K.A.M.), and (J.E.S.). (J.J.R.) and (L.A.B.) developed and characterized AST1825 cell line. (M.A.G) provided chemotherapeutic agents and pharmacology expertise. (L.A.B.) and (M.P.) wrote the manuscript, with all authors contributing to the writing and providing feedback.

## DATA AVAILABILITY

The RNA-SEQ dataset analyzed in this study is publicly available in the Gene Expression Omnibus (GSE89307). All data generated and supporting the findings of this study are available in the paper or are available from the corresponding author on request.

---

**Supplemental Figure 1.**
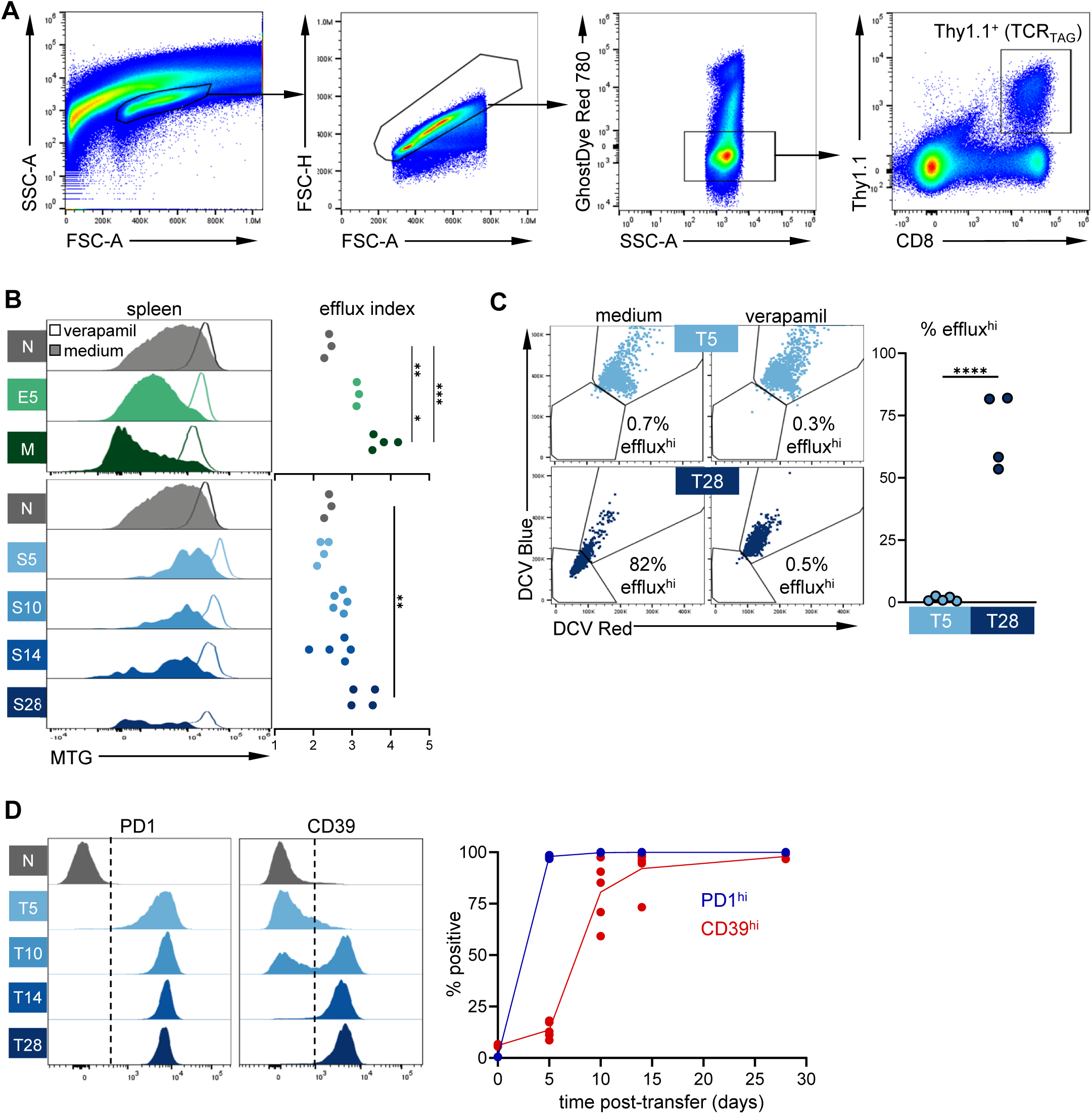
Gating strategy and additional characterization of efflux capacity and immunophenotype in liver and splenic TST. (A) Representative flow cytometry gating strategy for identification of TCR_TAG_ (Thy1.1^+^). This strategy is used in all adoptive transfer and *in vitro* experiments unless otherwise specified. **(B)** MTG fluorescence of TCR_TAG_ isolated from the spleens of infected or tumor-bearing mice as in **Fig. 1B**. Summary plot shows efflux index. \*\**P* ≤ 0.01, \*\*\**P*≤ 0.001; one-way ANOVA followed by Tukey’s multiple comparisons test, n = 3-5 per time point, representative of three independent experiments. (**C)** Dye cycle violet (DCV) staining of TCR_TAG_ at 5 and 28 days post-transfer as in **Fig. 1B** with gate set based on verapamil-treated control. Summary plot shows % efflux^hi^. \*\*\*\**P* ≤ 0.0001; two-tailed Student’s t-test, n = 4-5 per timepoint, representative of two independent experiments. **(D)** PD1 and CD39 expression of tumor TCR_TAG_ at indicated timepoints. Summary plot shows %PD1^hi^ and %CD39^hi^, with negative gates set on naive. n = 3-5 per timepoint, data representative of three independent experiments.

**Supplemental Figure 2.**
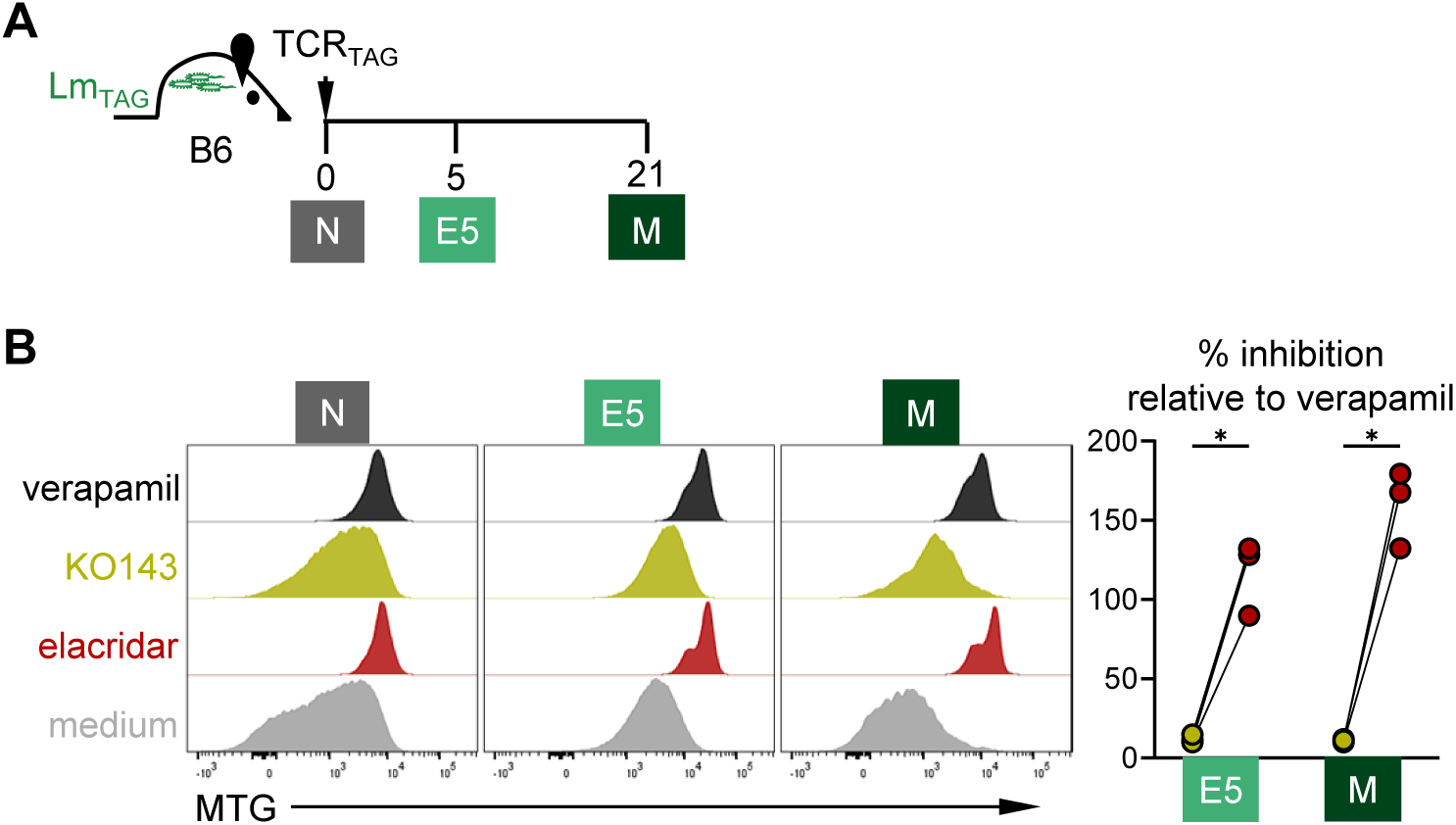
Effect of specific efflux inhibitors on effector and memory CD8 T cell efflux. (A) Experimental scheme: naive TCR_TAG_ were adoptively transferred into Lm_TAG_-infected B6 mice. Lymphocytes were re-isolated from livers and analyzed by flow cytometry at days 5 (effector) and 21 (memory) post-transfer. **(B)** MTG fluorescence in TCR_TAG_ with indicated inhibitors. Summary plot shows % inhibition calculated from MTG MFI compared to verapamil-treated control. \**P* ≤ 0.05; paired Student’s t-test with Holm-Šídák correction for multiple comparisons, n = 3 per group.

**Supplemental Table 1.**
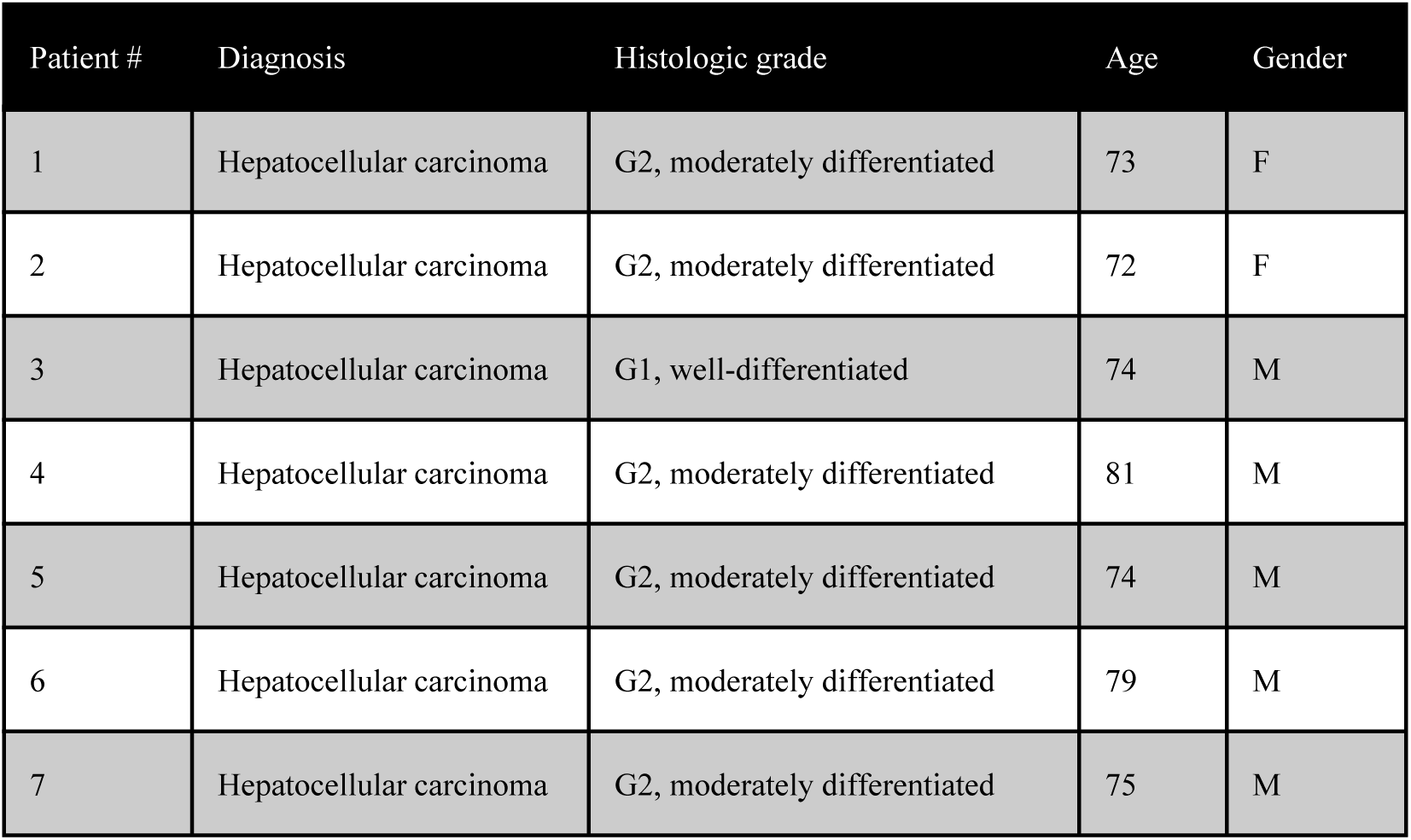
Patient characteristics.

